# Evidence for the Modulation of the Immune Response in Peripheral Blood Mononuclear Cells after Stimulation with a High Molecular Weight β-glucan Isolated from *Lactobacillus fermentum* Lf2

**DOI:** 10.1101/400267

**Authors:** Ana Vitlic, Sohaib Sadiq, Hafiz I. Ahmed, Elisa C. Ale, Ana G. Binetti, Andrew Collett, Paul N. Humpreys, Andrew P. Laws

## Abstract

*Lactobacillus fermentum* Lf 2 produces large amounts of exopolysaccharides under optimized conditions (∼2 g/L, EPS) which have been shown to possess immunomodulatory activity. In this study, the crude EPS was fractionated to give a high molecular weight (HMw) homoglycan and a mixture of medium molecular weight heteroglycans. The HMw EPS was isolated and identified as a β-glucan.

Peripheral blood mononuclear cells (PBMC) were pre-treated with purified polysaccharide to determine if the HMw β-glucan is responsible for the immunomodulatory activity. Cells were also stimulated with either lipopolysaccharide (LPS) or phytohemagglutinin (PHA) and their effects, both with and without β-glucan pre-treatment, compared.

Exposure of the cells to β-glucan increased their metabolic activity and whilst a small but statistically significant drop in CD14 expression was observed at Day 1, the levels were significantly elevated at Day 2. High levels of CD14 expression were observed in cells initially exposed to the β-glucan and subsequently stimulated with either LPS or PHA. In contrast, reduced levels of TLR-2 expression were observed for cells initially exposed to the β-glucan and subsequently stimulated with LPS.

TNF-α levels were elevated in β-glucan treated cells (Day1) with the levels dropping back once the β-glucan had been removed (Day 2). The stimulants LPS and PHA both induced significant rises in TNF-α levels, however, this induction was completely (LPS) or partially blocked (PHA) in β-glucan pre-treated cells.

The results indicate a role for the bacterial β-glucan in modulating the immune response following exposure to agonists such as bacterial LPS.

## IMPORTANCE

The use of foods containing probiotics (i.e. live microorganisms providing health benefits) is growing in popularity. Whilst there is empirical evidence identifying specific microorganisms as having beneficial effects, the precise mechanisms by which these strains exert influence is not well understood. One of the problems is that there are a large number of structures presented at the surface of microorganisms which can interact with the host and which can initiate a host response. In the current work, we addressed this issue by choosing a potential probiotic strain of bacteria associated with cheese production, *L. fermentum* Lf2, which produces large quantities of polysaccharides at its surface. From this polysaccharide mixture, we have isolated, purified and characterised a HMw β-glucan and demonstrated that it has immunomodulatory activity. This study demonstrates that the β-glucan produced by *L. fermentum* Lf2 can modify the immune response of PBMC cells when subjected to a bacterial endotoxin.

### 1. Introduction

Species of *Lactobacillus fermentum* are obligate heterofermentative Gram-positive bacteria that can be isolated from a wide variety of sources and include members that are normal inhabitants of the human microflora (1; 2). A significant number of strains are closely associated with the production of fermented foods (3-7) and a number have been exploited for their probiotic properties (8; 9). *L. fermentum* Lf2 is a strain that was isolated as a contaminant culture during cheese manufacture (10). The strain produces significant quantities of a mixture of EPS (∼2g/L) and it has been demonstrated that the crude EPS has immunomodulatory activity (11). In previous studies, using EPS from lactic acid bacteria (12; 13), we have shown that specific EPS can stimulate production of pro- and anti-inflammatory cytokines. We also provided evidence that the EPS can modulate the immune response and in doing so provide a degree of immunotolerance. It is likely that immunotolerance has evolved as a way of preventing damage that can occur through too robust an immune response and as a mechanism to prevent reaction against non-self but beneficial antigens, such as those produced by commensal microorganisms. Endotoxin tolerance is the ability of an organism to, if previously primed with a bacterial antigen, survive what would under normal circumstances be lethal doses of bacterial infection (14). While this temporary state of unresponsiveness to endotoxin can be detrimental (15) for example in sepsis, equally it can have a positive impact in pathologies that are the consequence of hyperactive immune response such as in inflammatory bowel diseases.

One of the potential mechanisms for inducing immunotolerance is by interaction of EPS with pattern recognition receptors, CD14 and TLR-2, present on the cell surface that are involved in recognising bacterial components including bacterial lipids (16; 17), lipoproteins, peptidoglycans (18) and, relevant to this study, EPS (19). In peripheral blood mononuclear cells (PBMC) CD14 and TLR-2 are present predominantly on monocytes and have an important role in innate host defence and they trigger activation of adaptive immunity. It has been shown that tolerance to bacterial lipoproteins is accompanied by down-regulation of TLR-2 expression (20). However, the role of CD14 in immune tolerance is more complex, with reports showing that development of a tolerant phenotype precedes CD14 down-regulation (21) thus demonstrating a CD14-independent mechanism of tolerance (22; 23).

The aim of the current study was to investigate the immunomodulatory activity of the EPS produced by *L. fermentum* Lf2 and more specifically to examine the effect of purified EPS on PBMC. These cells are easily isolated from whole blood and although there are some small differences in the composition and activation status compared with the immune cells of the gastrointestinal tract, they share certain key phenotypic characteristics (24) and have previously been used to study behaviour of mucosal immunity of the GIT (25). PBMC were isolated and pre-treated with different concentration of a purified HMw EPS from *L. fermentum* Lf2. The immunomodulatory activity was evaluated in these cultures as well as those that were subsequently stimulated with known activators of human blood cells such as lipopolysaccharide (LPS) from *E. coli* and the lectin phytohemagglutinin (PHA). The effect was measured by monitoring changes in the expression of the receptors involved in the interaction of bacteria and immune cells such as CD14 and TLR-2, and secretion of TNF-α, a known proinflammatory cytokine important in monocyte and T cell signalling.

## RESULTS

### Composition of the crude exopolysaccharide (EPS) and purification of the High Molecular weight exopolysaccharide (HMw EPS)

SEC-MALLS analysis of the crude EPS produced by *L. fermentum* Lf2 identified a complex mixture of biomacromolecules. The three most significant peaks included a HMw EPS (RT= 25 min), a mixture of medium molecular weight polysaccharides (average RT= 35 min) and a UV absorbing species (RT= 40 min) (Fig 1A). After preparative size exclusion chromatography a pure sample of the HMw EPS was obtained in fractions 10-19. The combined fractions were concentrated, freeze-dried and redissolved in deuterium oxide (D_2_O) before SEC-MALLS analysis. The SEC-MALLS trace (Fig 1B) contained a single peak (RT= 26 mins) with a weight average molecular mass of 1.23 × 10^6^ Da and a polydispersity value of 1.104.

**Figure 1.**
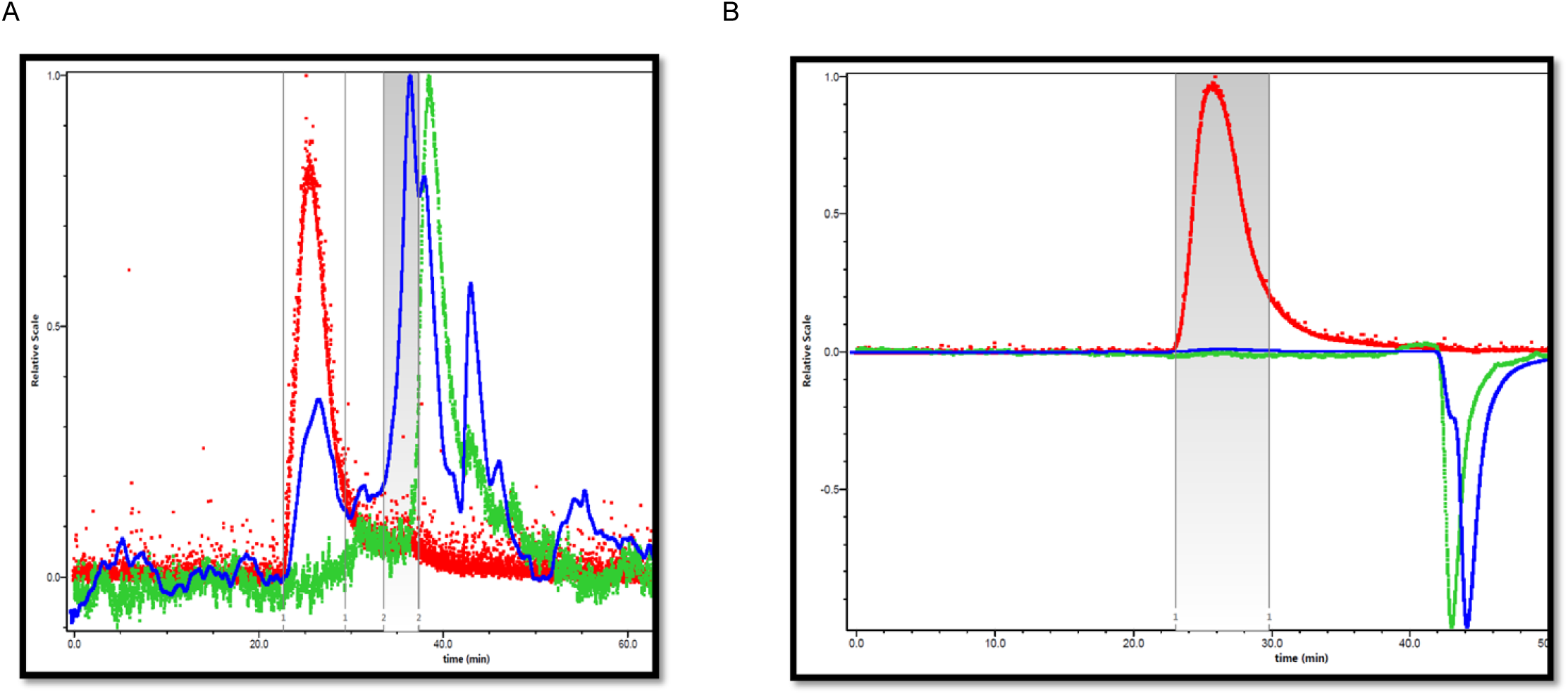
Analytical SEC-MALLS traces; Red trace = molecular weight dependent light scattering, Blue trace = concentration dependent refractive index, Green trace= UV absorption at 260 nm A) Chromatograph for the crude EPS from *Lactobacillus fermentum* Lf2 ((1 mg/mL) in aq. NaNO_3_ (0.1M). B) Chromatograph for the HMw EPS from *Lactobacillus fermentum* Lf2 ((1 mg/mL) in aq. NaNO_3_ (0.1M).

### Characterization of the HMw EPS

Monomer analysis and determination of the absolute configuration of the monosaccharides identified that the HMw EPS was composed entirely of D-glucopyranose. Linkage analysis, using permethylated alditol acetates, confirmed the presence of a 1,5-di-*O*-acetyl-2,3,4,6-tetra-*O*-methylglucitol (corresponding to a terminal glucose), a 1,3,5-tri-*O*-acetyl-2,4,6-tri-*O*-methylglucitol (corresponding to a 1,3-linked glucose) and a 1,2,3,5-tetra-*O*-acetyl-4,6-di-*O*-methylglucitol (corresponding to a 1,2,3-linked glucose).

The ^1^H-NMR spectrum (Fig 2) had two distinct resonances in the anomeric region (4.93 & 4.85 ppm) with an integral ratio very close to 1:2 and both had large ^3^*J*_H1,H2_-coupling constants which are indicative of β-linked hexoses.

**Figure 2.**
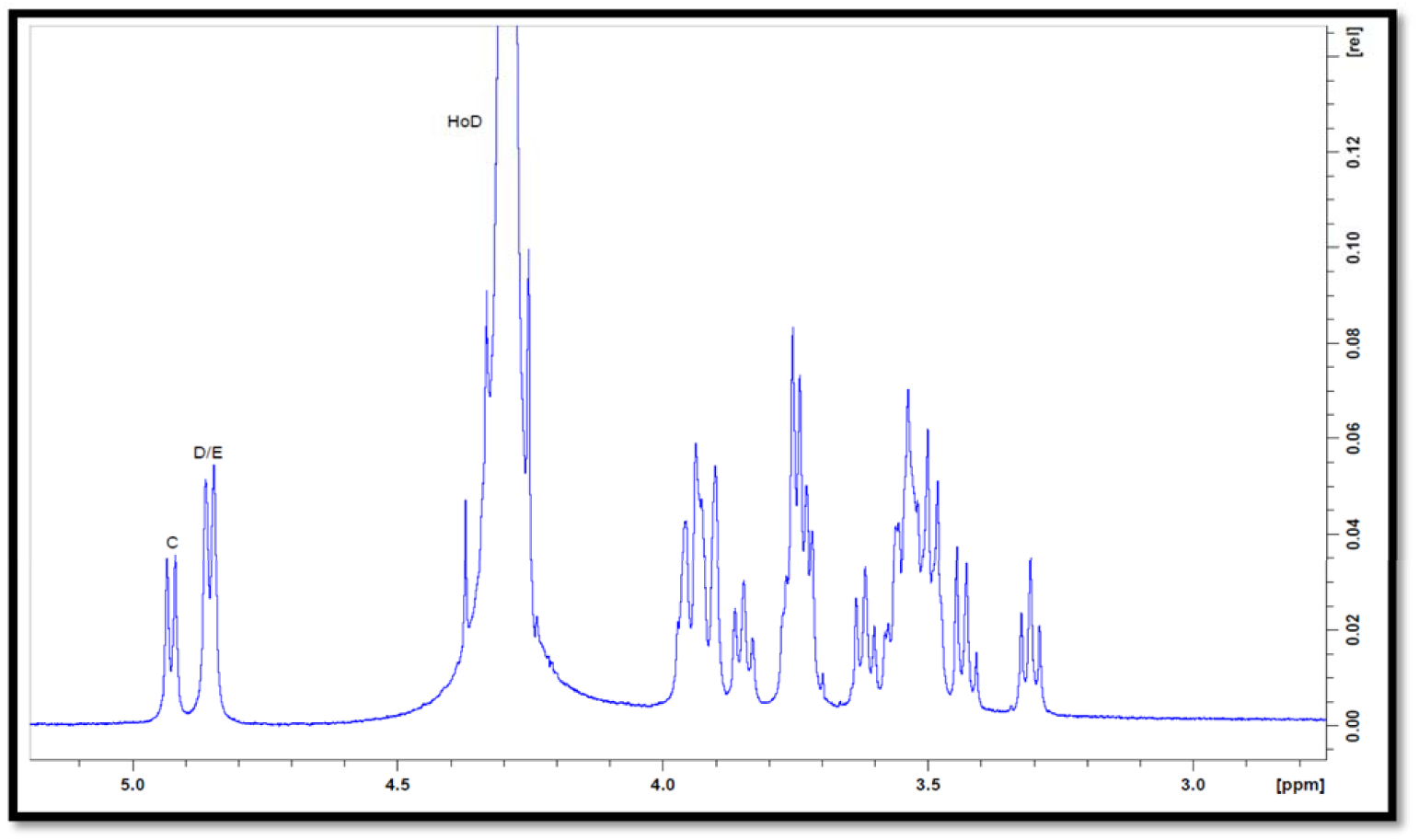
^1^H-NMR for the HMw EPS recorded in solution in D_2_O (5-10 mg in 0.65 mL) at 70 °C on a Bruker 500MHz spectrometer. The residual HOD signal is at 4.29 ppm at 70 ^°C^.

The location of the ring protons was identified through inspection of a combination of ^1^H-^1^H-COSY and ^1^H-^1^H-TOCSY spectra (see supplementary information) and that of the ring carbons using a combination of a ^1^H-^13^C-HSQC-spectrum and a ^1^H-^13^C-HSQC-TOCSY spectrum (see supplementary information). The chemical shifts for the proton and carbons are listed in Table 1.

**Table 1.** Proton and carbon chemical shifts (ppm) for the Lf2 HMW-EPS recorded in D_2_O at 70 °C on a Bruker Avance 500MHz NMR spectrometer. Chemical shifts are reference to external acetone 2.225 ppm for ^1^H and 31.55 ppm for ^13^C

|  | H-1<br>C-1 | H-2<br>C-2 | H-3<br>C-3 | H-4<br>C-4 | H-5<br>C-5 | H-6a<br>C-6a | H-6b<br>C6b |
| --- | --- | --- | --- | --- | --- | --- | --- |
| A | 4.93<br>102.7 | 3.31<br>74.7 | 3.50<br>76.3 | 3.50<br>76.3 | 3.50<br>76.3 | 3.96<br>61.6 | 3.75<br>61.6 |
| B | 4.85<br>102.2 | 3.84<br>80.1 | 3.95<br>84.2 | 3.55<br>68.8 | 3.55<br>68.8 | 3.94<br>61.6 | 3.74<br>61.4 |
| C | 4.85<br>102.2 | 3.62<br>73.2 | 3.75<br>86.6 | 3.42<br>70.6 | 3.42<br>70.6 | 3.90<br>61.6 | 3.73<br>61.4 |

Finally, the linkage pattern of the monosaccharides in the repeat unit was confirmed by examination of a ROESY and a HMBC spectrum and identification of the inter-residue NOEs and long-range scalar coupling which were compared with data reported in the literature for identical EPS(26; 27) (see supplementary information). U.V. absorption and ^31^P NMR spectra were recorded for the HMw polysaccharide and these were not able to detect any phosphorous, protein or nucleic acid, and when taken along with the results of the other characterization studies, this confirmed that a high purity polysaccharide had been isolated.

### PBMC isolation and cell culture

In the Day 1 experiment PBMC were activated by treatment with the HMw β-glucan in all the donors (p < 0.001 - 0.0001) (Fig 3). The levels of activation, compared to the control, were comparable for both β-glucan concentrations used (50 µg/mL and 100 µg/mL), except for Donor B where higher levels were obtained for 50 µg/mL EPS (p < 0.0001).

In the Day 2 experiment, levels of activation remained elevated in the β-glucan treated conditions for two donors, regardless of the type of subsequent stimulation (p < 0.05-0.0001). Treatment with LPS alone also showed higher metabolic activity as measured by the MTS assay, and this was true regardless of whether or not they had been pre-treated with β-glucan (p < 0.0001).

**Figure 3.**
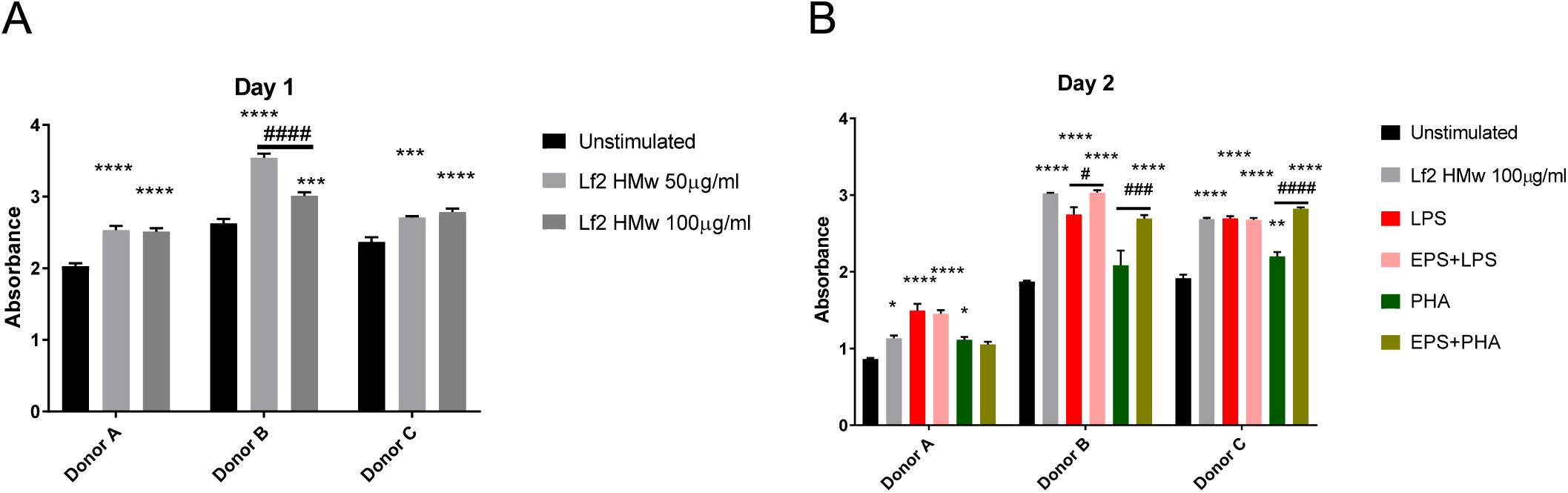
Non-radioactive MTS assay was used for measuring metabolic activity of the cells under different culture conditions as measured after A. Experiment 1, Day 1 and B. Experiment 2, Day 2. For Experiment 1, Unstimulated conditions were left untreated while Lf2 HMw 50µg/ml and Lf2 HMw 100µg/ml were treated with EPS at indicated concentrations on Day 0 and incubated at 37°C, 5% CO_2_. Following 24h incubation, cells were harvested and 1×10^5^ cells/well were transferred into 96-well plates for MTS assay performed following the supplier’s instructions. For Experiment 2, Day 2, Unstimulated condition was left untreated on Day 0 then washed and left unstimulated on Day 1. Lf2 HMw 100µg/ml were pre-treated at Day 0 with indicated concentration of EPS, then washed and left unstimulated on Day 1. LPS and PHA alone are conditions that were left untreated on Day 0, then washed and subsequently stimulated with 100ng/ml LPS and 10µg/ml PHA, respectively. EPS + LPS and EPS + PHA represent the conditions that were pre-treated with EPS Lf2 HMw on Day 0, washed and stimulated with 100ng/ml LPS and 10µg/ml PHA, respectively on Day 1. Following 24h incubation, cells were harvested and 1×10^5^ cells/well were transferred into 96-well plates for MTS assay performed following the supplier’s instructions. Two-Way ANOVA was used to compare the conditions for each donor. Data are presented as Mean ± SEM. * p < 0.05, ** p < 0.01, *** p < 0.001, **** p < 0.0001 when compared to the untreated, unstimulated condition; # p < 0.05, ### p < 0.001, #### p < 0.0001 when compared between two indicated conditions.

Apparent anomalies in the MTS results were observed for PHA treatment, where although a statistically significant activation was observed for two out of three donors tested (p < 0.05-0.01), the recorded levels did not correspond to the level of cell proliferation observed on microscopic images for this condition (Fig 4). It is possible that the conditions used for PHA stimulation interfered with the results of the MTS assay such that the level of metabolic activity that this assay measured did not accurately correspond to the level of proliferation exhibited after PHA stimulation (28; 29).

**Figure 4.**
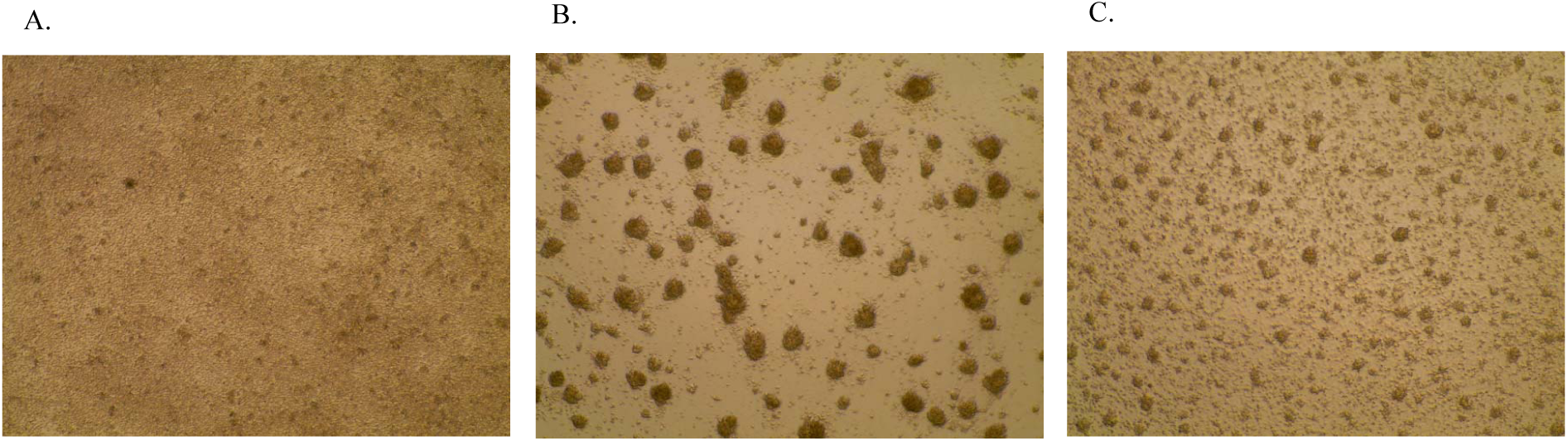
Representative microscopic images showing the effect of PHA with and without EPS pre-treatment on PBMC proliferation as seen at Day 2 of experiment 2. A. Untreated at Day 0, then washed and unstimulated at Day 1; B. Untreated at Day 0, then washed and stimulated with 10µg/ml PHA at Day 1; C. Pre-treated with 100µg/ml Lf2 HMw EPS at Day 0, then washed and stimulated with 10µg/ml PHA at Day 1.

Microscopic inspection of the cells (Fig 4) confirmed the high levels of proliferation and clumping of cells after PHA stimulation when compared to the unstimulated and untreated condition (Fig 4A) and the degree of cell aggregation was attenuated by previous treatment with β-glucan (Fig 4C). Absorbance values measured by MTS assay confirmed the difference between PHA only stimulated condition compared to the EPS pre-treated and PHA stimulated condition, at least in Donors B and C, however, the levels were lower than what would be expected for the higher proliferation visible in the images of PHA stimulated PBMC.

### Flow cytometric assessment of pattern recognition receptors CD14 and TLR-2

Changes in expression, presented as median fluorescence intensity (MFI), of the CD14 and TLR-2 receptors that are involved in recognition of microbial-specific molecules are presented in Figures 5 and 6, respectively. In the Day 1 experiment, for two out of three donors CD14 showed lower expression after pre-treatment with EPS compared to the untreated condition, regardless of the concentration used (p < 0.0001, Fig 5). In the second experiment, at Day 2, however, initial pre-treatment with EPS led to a robust increase in the CD14 expression, for all conditions, compared to those that had not been pre-treated with EPS (untreated unstimulated vs EPS pre-treated; LPS stimulated vs EPS pre-treated and LPS stimulated, with p < 0.0001 in all cases, and in Donor B only, PHA stimulated vs EPS pre-treated and PHA stimulated, p < 0.01; Fig 5B). LPS stimulation led to a moderate decrease in CD 14 expression in two out of three donors (p < 0.0001 for Donor B and p < 0.01 for Donor C), while the levels of CD14 markers were higher in Donor A (p < 0.0001), when compared to the untreated and unstimulated condition at Day 2.

Unsurprisingly, PHA consistently attenuated CD14 expression (p <0.0001 for all three donors), while pre-treatment with EPS marginally reduced the decrease in expression caused by PHA, as seen in Donor B (p < 0.01).

**Figure 5.**
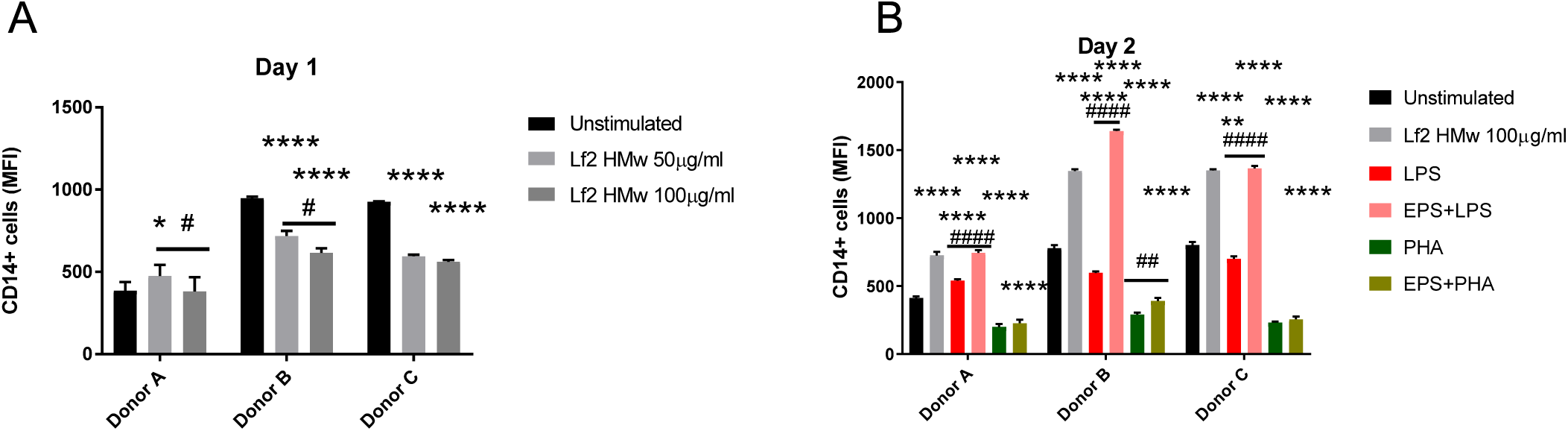
Changes in CD14 expression on PBMC following different conditions presented as median fluorescence intensity (MFI) of CD14^+^ cells after A. Day 1 and B. Day 2 experiment. For Day 1, Unstimulated conditions were left untreated while Lf2 HMw 50µg/ml and Lf2 HMw 100µg/ml were treated with EPS at indicated concentrations on Day 0 and incubated at 37°C, 5% CO_2_. Following 24h incubation, cells were harvested and stained with anti-CD14 antibody and analysed by flow cytometry. For Day 2 experiment, Unstimulated condition was left untreated on Day 0, then washed and left unstimulated on Day 1. Lf2 HMw 100µg/ml were pre-treated at Day 0 with indicated concentration of EPS, then washed and left unstimulated on Day 1. LPS and PHA alone are conditions that were left untreated on Day 0, then washed and subsequently stimulated with 100ng/ml LPS and 10µg/ml PHA, respectively. EPS + LPS and EPS + PHA represent the conditions that were pre-treated with EPS Lf2 HMw on Day 0, then washed and stimulated with 100ng/ml LPS and 10µg/ml PHA, respectively on Day 1. Following 24h incubation, cells were harvested and stained with anti-CD14 antibody and analysed by flow cytometry. Two-Way ANOVA was used to compare the conditions for each donor. Data are presented as Mean ± SEM. * p < 0.05, *** p < 0.001, **** p < 0.0001 when compared to the untreated, unstimulated condition; # p < 0.05, ## p < 0.01, #### p < 0.0001 when compared between two indicated conditions. Gates were set on single cells and debris was excluded using FSC vs SSC plots. Gating was performed using FMO (Fluorescence Minus One) controls.

The effect of PBMC stimulation on TLR-2 expression was less consistent and the only result of note is that there was a clear decrease at Day 2 in TLR-2 expression after EPS pre-treatment, followed by LPS stimulation, compared to the LPS stimulated alone (p< 0.05 for Donor C and p < 0.0001 for Donors A and B, Fig 6).

**Figure 6.**
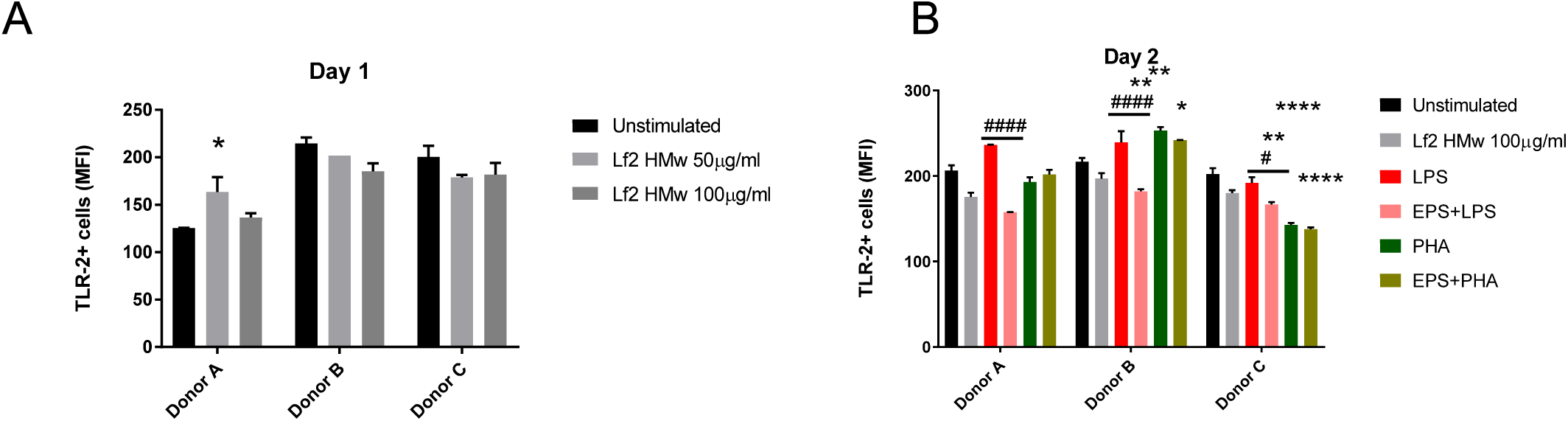
Changes in TLR-2 expression on PBMC following different conditions presented as median fluorescence intensity (MFI) of TLR-2^+^ cells after A. Day 1 and B. Day 2 experiment. For Day 1, Unstimulated conditions were left untreated while Lf2 HMw 50µg/ml and Lf2 HMw 100µg/ml were treated with EPS at indicated concentrations on Day 0 and incubated at 37°C, 5% CO_2_. Following 24h incubation, cells were harvested and stained with anti-TLR-2 antibody and analysed by flow cytometry. For Day 2 experiment, Unstimulated condition was left untreated on Day 0, then washed and left unstimulated on Day 1. Lf2 HMw 100µg/ml were pre-treated at Day 0 with indicated concentration of EPS, then washed and left unstimulated on Day 1. LPS and PHA alone are conditions that were left untreated on Day 0, then washed and subsequently stimulated with 100ng/ml LPS and 10µg/ml PHA, respectively. EPS + LPS and EPS + PHA represent the conditions that were pre-treated with EPS Lf2 HMw on Day 0, then washed and stimulated with 100ng/ml LPS and 10µg/ml PHA, respectively on Day 1. Following 24h incubation, cells were harvested and stained with anti-TLR-2 antibody and analysed by flow cytometry. Two-Way ANOVA was used to compare the conditions for each donor. Data are presented as Mean ± SEM. * p < 0.05, ** p < 0.01, *** p < 0.001, **** p < 0.0001 when compared to the untreated, unstimulated condition; # p < 0.05, #### p < 0.0001 when compared between two indicated conditions. Gates were set on single cells and debris was excluded using FSC vs SSC plots. Gating was performed using FMO (Fluorescence Minus One) controls.

### Stimulation of TNF-α secretion

Initial treatment with EPS resulted in an increase in TNF-α secretion on Day 1 for both concentrations (Fig 7A). However, while in Donors B and C the increase in TNF-α was greater for the lower EPS concentration (50 µg/ml), the opposite was true for Donor A (Fig 7A).

Interestingly, TNF-α levels in the Day 2 experiment for the cells that were pre-treated with EPS at Day 0 and then left unstimulated at Day 1 were comparable to control cells (Fig 7B) indicating a robust decrease in the levels of this cytokine when compared to EPS pre-treated conditions in the Day 1 experiment (Fig 7A). For cells that were treated with EPS on Day 0 and then stimulated on Day 1 with LPS or PHA the pre-treatment with EPS greatly attenuated a robust increase in TNF-α secretion for all donors (p < 0.01 between LPS and EPS+LPS in Donor B, and p < 0.0001 between PHA and EPS+PHA in Donors A and C).

**Figure 7.**
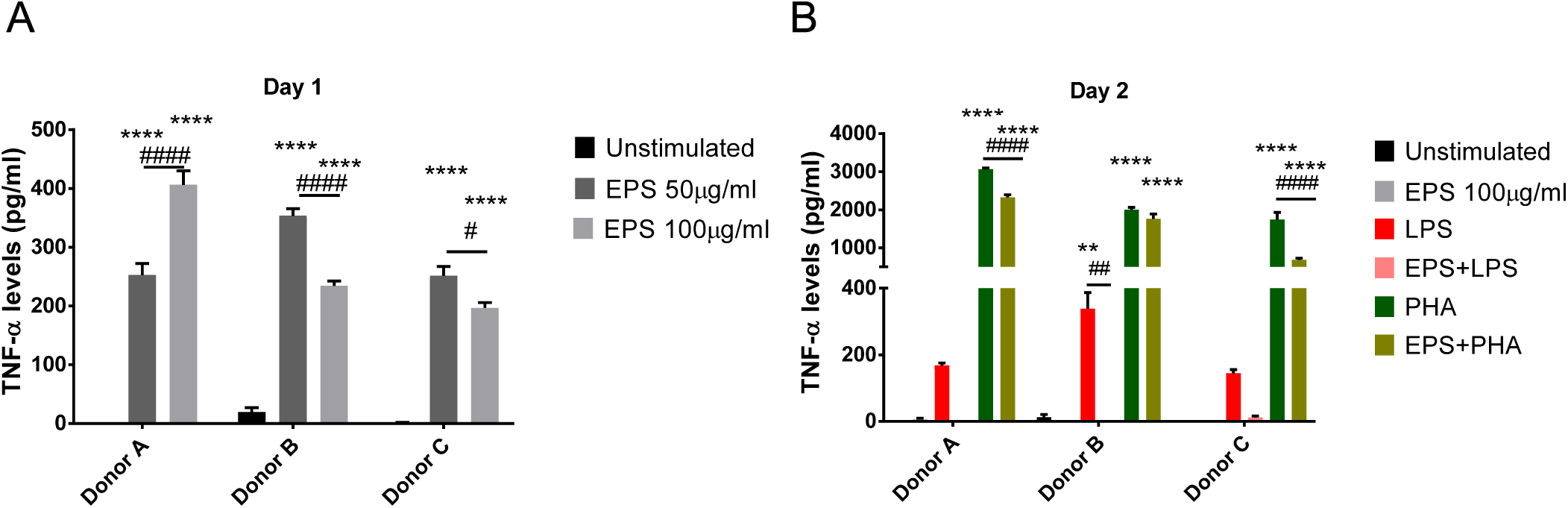
Changes in TNF-α levels in cell culture supernatants following different conditions after A. Day 1 and B. Day 2 experiment. For Day 1, Unstimulated conditions were left untreated while Lf2 HMw 50µg/ml and Lf2 HMw 100µg/ml were treated with EPS at indicated concentrations on Day 0 and incubated at 37°C, 5% CO_2_. Following 24h incubation, supernatants were stored at −80°C until subsequent analysis for TNF-α levels by ELISA. For Day 2 experiment, Unstimulated condition was left untreated on Day 0, then washed and left unstimulated on Day 1. Lf2 HMw 100µg/ml were pre-treated at Day 0 with indicated concentration of EPS, then washed and left unstimulated on Day 1. LPS and PHA alone are conditions that were left untreated on Day 0, then washed and subsequently stimulated with 100ng/ml LPS and 10µg/ml PHA, respectively. EPS + LPS and EPS + PHA represent the conditions that were pre-treated with EPS Lf2 HMw on Day 0, then washed and stimulated with 100ng/ml LPS and 10µg/ml PHA, respectively on Day 1. Following 24h incubation, supernatants were stored at −80°C until subsequent analysis for TNF-α levels by ELISA. Two-Way ANOVA was used to compare the conditions for each donor. Data are presented as Mean ± SEM. ** p < 0.01, **** p < 0.0001 when compared to the untreated, unstimulated condition; # p < 0.05, ## p < 0.01, #### p < 0.0001 when compared between two indicated conditions.

## DISCUSSION

The aim of this study was determine which component of the bacterial EPS produced by *L. fermentum* Lf2 is responsible for the immunomodulatory properties of the crude EPS and to gain an understanding of the nature of this modulation. Analysis indicated that the crude EPS included both a HMw polysaccharide and a mixture of medium molecular weight polysaccharides that were separated by size exclusion chromatography. The HMw polysaccharide was identified as a β-glucan with a weight average molecular mass (1.23 × 10^6^ Da) which is similar to that of EPS obtained from other strains of lactic acid bacteria (LAB) (30; 31). Using NMR analysis, the ^1^H and ^13^C-chemical shifts were recorded for the β-glucan (Table 1) these are identical to those published for the EPS isolated from *Pediococcus damnosus* 2.6 (26), *Oenococcus oeni* I4 (32) and for one of the two EPSs produced by *Lactobacillus sp. G-77*(27). The results of the characterisation studies are consistent with the β-glucan having the following trisaccharide repeating unit:

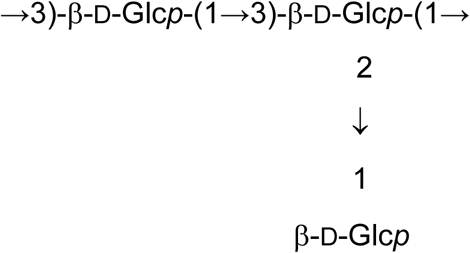

It is worth noting that the repeating unit is similar to the type 37 polysaccharide produced by *Streptococcus pneumoniae* which has the same linear backbone (→3)- *β*-D-Glc*p*-(1→) bearing a terminal glucose unit at the 2-position but these appear on each residue in the *S. pneumoniae* EPS, rather than on alternate residues as in the *L. fermentum* Lf2 β-glucan (33). All attempts to purify the medium molecular weight polysaccharide were unsuccessful, the on-going work to characterise this material will be reported elsewhere.

Treatment of PBMC with the β-glucan had no impact on cell viability but did increase the metabolic activity of cells that was measured as an increase in mitochondrial dehydrogenase activity. At the concentrations employed, the β-glucan was able to activate PBMC in culture compared to controls at Day 1, indicating a functional interaction between the two. Interestingly, this activation persisted for 24 h after the β-glucan had been removed from solution. This is in agreement with previous reports where cell wall extracts and purified EPS stimulated cell proliferation of different immune cell types: including BALB/c mice splenocytes and mesenteric lymphocytes (34) using EPS from LAB, and mouse lymphocytes by EPS from *Paenibacillus jamilae* CP-7 (35). The stimulatory effect of EPS has also been shown to be concentration dependent when two *Lactobacillus rhamnosus* strains (ATCC 9595 and RW-9595M), that differ in the amount of EPS they produce, were used to stimulate splenic lymphocytes (36). In contrast, EPS from bifidobacteria has been reported to have anti-proliferative activity on rat PBMC and gut associated lymphoid tissue cells (37). Whilst other authors have reported bifidobacteria EPS stimulates proliferation of the intestinal Caco-2 but not HT-29 (38) cells while the EPS from *Lactobacillus gasseri* strains G10 and H15 had no effect on the proliferation of the cervical cancer cell line HeLa (39). These differences are not surprising since it has previously been demonstrated that EPS can exert a differential effect on the metabolic activity of different cell types (40).

The MTS assays gave results in agreement with previous reports, where the optimal PHA concentration for cell stimulation was 2.5µg and where only modest increase in metabolic activity was reported for cultures stimulated with 10µg/ml PHA (28). From inspection of the microscope images it is clear that pre-treatment with the β-glucan partially reversed the effect of PHA by reducing the amount of cell clumping.

Changes in cytokine production in response to the presence of microbial strains and their products is a way of demonstrating immunomodulatory effect on host cells. For example in our previous studies we demonstrated that the EPS obtained from two *Lactobacillus* strains, *L. helveticus ssp. rosyjski* and *L. acidophilus*, both increased IL-8 production in HT29-19A intestinal cell line (41). Other authors have reported changes in a number of pro-inflammatory cytokines, such as IL-12, IL-6 and TNF-α with pre-treatment with EPS from *L. rhamnosus* GG in combination with LPS stimulation when compared to the LPS stimulation alone (42). In the present study, pre-treatment with β-glucan stimulated TNF-α production after Day 1 with the levels falling back to those observed in the control 24 h after the β-glucan had been removed from solution.

One of the most significant results was observed for the cells which had been pre-treated with the β-glucan and which were subsequently stimulated by LPS or PHA, this led to large decreases in the levels of this cytokine, when compared to the corresponding stimulation alone (LPS and PHA, respectively). As TNF-α is a key proinflammatory cytokine, this is typical of an immune tolerant phenotype, with potentially important implications given its central role in the aetiology of the inflammatory bowel diseases such as Crohn’s disease and ulcerative colitis.

To better understand the interaction of the β-glucan with PBMC, expression of CD14 and TLR-2, receptors commonly involved in host microbial responses (43), was examined. An initial downregulation of CD14 on Day 1 (observed in two donors) suggests the involvement of this co-receptor in the β-glucan signalling. A similar small decrease in CD14 MFI on PBMC after 24 h stimulation with LPS has been previously reported (44), unsurprisingly since LPS is known to use CD14 as a co-receptor. Downregulation of CD14 after LPS stimulation was also observed in alveolar macrophages (23). A typical tolerance phenotype involves a permanent decrease in CD14 expression (23; 45). In contrast, in the present study, pre-treatment with β-glucan reversed any down-regulatory effect of LPS completely and, to a lesser extent, that of PHA. This positive influence on the availability of CD14 on PBMC could have potentially important positive effects: an endotoxin tolerant phenotype can have a detrimental effect on health and survival, such as with sepsis, where the risk of new infections is increased due to the refractory state of the organism (46). Thus the ability of the β-glucan to reverse a tolerance phenotype, at least in respect to CD14 expression, could be beneficial in diseases where continuous downregulation of CD14 reduces an organisms’ ability to respond to subsequent infections.

The influence of β-glucan treatment of PBMC alone on TLR-2 expression at Day 1 was not easy to determine, with both small increases and decreases in expression levels being observed. TLR-2 expression in cells pre-treated with β-glucan and subsequently stimulated with LPS decreased when compared to cells stimulated with LPS only. Similarly, it has been previously reported that treatment with *L. rhamnosus* in combination with LPS reduced TLR-2 expression at the mRNA level when compared to LPS stimulation alone (42). By contrast, an increase in cells expressing TLR-2 was observed when immune cells from mouse Peyer patches were stimulated with *Lactobacillus casei* (47). The range of effects that bacterial components exert on immune cells is not unexpected as this is known to be strain dependent (48). Modulation of TLR-2 by different microorganisms and their components in immunomodulatory phenotypes can cause changes in both pro- and anti-inflammatory cytokine secretion.

The data, in combination with recovery of CD14 expression and continuous activation of cells as demonstrated by the MTS assay on Day 2 suggest a difference in mechanism in which this β-glucan exerts tolerance when compared to classical endotoxin tolerance. It is therefore possible that the β-glucan could have a potentially important immunoprotective role. As such further investigations should address the mechanism of these effects and their *in vivo* relevance.

## MATERIAL AND METHODS

Unless otherwise stated, all reagents were purchased from Sigma-Aldrich Company Ltd. (Poole, Dorset UK) and were used as supplied.

### EPS production and purification

*L. fermentum* Lf2 was grown at the Instituto de Lactologia Industrial (Santa Fe, Argentina). The growth conditions, as well as the crude EPS extraction and purification methods have previously been reported (11; 49). Size Exclusion Chromatography coupled with Multi Angle Laser Light Scattering (SEC-MALLS-Wyatt technology, Santa Barbara, CA, USA) was used to determine the composition of the crude EPS. EPS samples (1 mg/mL) were prepared in aq. NaNO_3_ (0.1M) and stirred for 16 h to ensure the EPS was completely dissolved. Samples (100 µL) were injected in triplicate into a SEC-MALLS system (comprising of three columns connected in series: PL Aquagel-OH 40, 50 and 60 (8 μm, 30 cm x 7.5 mm, Agilent, Cheadle, UK) with a flow rate of 0.7 mL/min. A differential refractometer (Optilab rEX, Wyatt technology, Santa Barbara, CA, USA) was used to determine the concentration of the polysaccharide and a Dawn-EOS MALLS detector (laser operating at 690 nm) was used to determine the weight average molecular mass of the polysaccharide. An in-line UV detector (Shimadzu, Milton Keynes, UK) was used for the detection of proteins and nucleic acids. ASTRA version 6.0.1 software (Wyatt technology, Santa Barbara, CA, USA) was used for the data analysis.

The crude EPS was purified by preparative size exclusion chromatography on a Sephacryl S-500 HR column (XK26/60-GE Healthcare, Fisher Scientific, UK) eluting with ultrapure water at a flow rate of 5.0 mL min^-1^ using a FPLC kit (AKTA PRIME-Amersham Pharmacia, Biotech, GE Healthcare Life Sciences, Buckinghamshire, UK) with a set wavelength of 254 nm. In total, sixty 5 mL fractions were collected and the location of polysaccharides in the different fractions was identified by determining the carbohydrate content of each fraction using the Dubois method (50; 51). Fractions containing EPS were pooled and freeze-dried.

### NMR analysis of the high molecular weight EPS

NMR spectra of EPSs were recorded in solution in D_2_O (5-10 mg in 0.65 mL) and were run either at room temperature or at an elevated temperature (70 °C). All of the NMR spectra were recorded on a Bruker Avance 500.13 MHz spectrometer (Bruker-biospin, Coventry, UK) using Bruker’s TOPSPIN 4.0.1 software for analysis. Chemical shifts are expressed in ppm relative to internal acetone, 2.225 for ^1^H and 31.55 for ^13^C. A series of 2D-spectra were recorded including: a 2D gradient-selected double quantum filtered correlation spectrum (gs-DQF-COSY) recorded in magnitude mode at 70°C; a total correlation spectroscopy (TOCSY) experiment recorded with a mixing times of 120 ms; ^1^H-^13^C heteronuclear single quantum coherence (HSQC) spectra (decoupled and coupled); a heteronuclear multiple bond correlation (HMBC) spectrum; and finally, a rotating frame nuclear Overhauser effect spectrum (ROESY, mixing time of 200 ms). The 2D spectra were recorded with 256 experiments of 1024 data points. For the majority of spectra, time-domain data were multiplied by phase-shifted (squared-) sine-bell functions. After applying zero-filling and Fourier transformation, data sets of 1024-1024 points were obtained.

### Composition of the high molecular weight polysaccharide

The monosaccharides present were determined after acid hydrolysis either directly using HPAEC-PAD analysis or as their alditol acetates, as previously described(12). The absolute configuration of the sugars was determined by preparation of their respective 2-(*S*)-butylglycosides using Gerwig’s method(52). For linkage analysis, the samples were permethylated using the procedures described by Stellner(53).

### PBMC isolation and cell culture

Venous blood was collected after obtaining informed consent from three healthy volunteers in blood collecting tubes containing citrate-dextrose solution (ACD) at a whole blood to anticoagulant ratio 9:1. The study was approved by the Local Ethics Committee (School of Applied Sciences, University of Huddersfield, UK). Blood samples were layered over Histopaque^®^-1077 and PBMC isolated using a standard density gradient centrifugation protocol. Enriched PBMC were suspended in RPMI 1640 medium (Gibco, Life Technologies, UK) supplemented with 4 mM L-Glutamine, 50 IU/ml/50 μg/ml/ Penicillin/Streptomycin and 20 mM HEPES (all from Gibco, Life Technologies, UK) and 10% heat-inactivated fetal bovine serum (FBS) and 2×10^6^ cells/well were added to 24-well plates (Starlab, UK). The cells were then challenged as shown in Figure 8. For the Day 1 experiment (Fig 8A), cells were treated with two different concentrations of EPS (50 µg/mL and 100µg/mL) or left untreated, four replicates were set up for each condition. For the Day 2 experiment (Fig 8B), at Day 0, half of the wells were treated with 100 µg/mL EPS while the other half was left untreated. Cells were incubated for 24 h, at 37 °C, 5% CO_2_ after which they were washed with supplemented RPMI 1640 and stimulated with either LPS (Lipopolysaccharides from *Escherichia coli* O111:B4) at a final concentration of 100 ng/mL or PHA-M (Lectin from *Phaseolus vulgaris* (red kidney bean)) at 10 µg/ml, or left unstimulated (Fig 8B). (final incubation of 24 h, at 37 °C, 5% CO_2_)Each condition was set up in four replicates.

**Figure 8.**
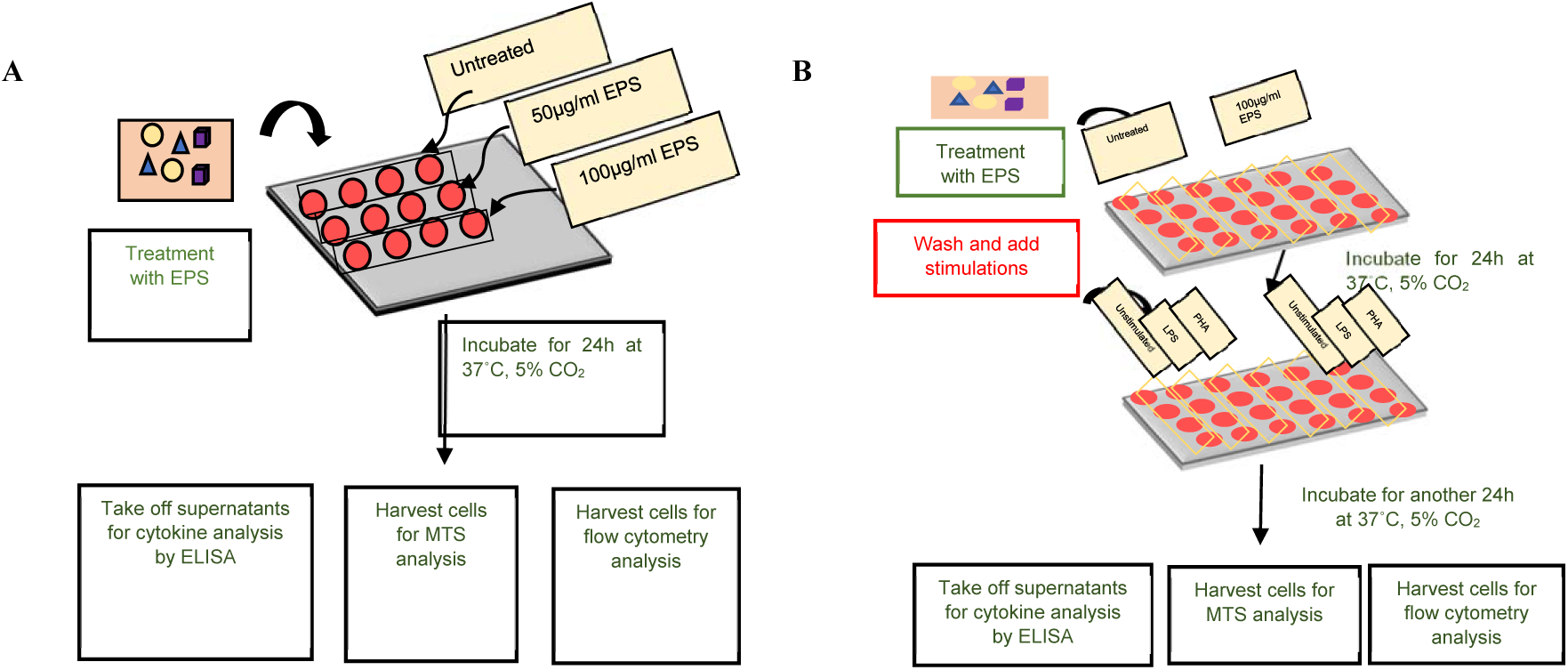
Schematic representation of the cell culture protocol completed on A. Experimet 1 and B. Experiment 2. A. On Day 0, cells were treated with two different Lf2 HMw EPS concentrations (50µg/ml and 100µg/ml) or left untreated at 37°C, 5% CO_2_. Following 24h incubation, supernatants were taken off and stored at −80°C for subsequent analysis by ELISA. Cells were harvested and 1×10^5^ cells/well were transferred into 96-well plates for MTS assay performed following the supplier’s instructions. The remaining cells were stained with antibodies for CD14 and TLR-2 and analysed for the expression of these markers by flow cytometry. B. on Day 0 cells were pre-treated with 100µg/ml EPS or left untreated and incubated for 24h at 37°C, 5% CO_2_. On Day 1, cells were washed to remove EPS and were stimulated with LPS (100ng/ml) or PHA (10µg/ml) or left unstimulated and incubated for another 24h at 37°C, 5% CO_2._ On Day 2, supernatants were then stored at −80°C for subsequent cytokine analysis by ELISA. As before 1×10^5^ of harvested cells/well were transferred into 96-well plate for MTS assay while remaining cells were stained for flow cytometric analysis of CD14 and TLR-2 expression.

### MTS assay

In order to measure metabolic activity of the cell cultures under different conditions, the CellTiter 96^®^ AQueous One Solution Cell Proliferation Assay, MTS assay (Promega, UK) was used as per the manufacturer’s instructions. At the end of each experiment 1×10^5^ cells from individual wells were transferred into separate wells of a 96-well plate and tetrazolium compound [3-(4,5-dimethylthiazol-2-yl)-5-(3-carboxymethoxyphenyl)-2-(4-sulfophenyl)-2H-tetrazolium, inner salt; MTS) and an electron coupling reagent (phenazine ethosulfate; PES) were added and the absorbance at 490nm was measured.

### Flow cytometry

To monitor CD14 and TLR-2 expression the remaining PBMC were washed with FACS buffer (PBS, Lonza, UK, 1% Bovine serum albumin, BSA, 0.1% Sodium azide, Sigma Aldrich, UK), and fixed with 1% paraformaldehyde (PFA, Sigma Aldrich, UK). Cells were then washed with permeabilisation solution (Perm Wash, BD Biosciences, UK) and stained with anti-CD14 and anti-TLR-2 antibodies (Biolegend, UK). Following the incubation, cells were washed with Perm Wash buffer and resuspended in 1% PFA until acquisition using The Guava^®^ easyCyte flow cytometer (Merck Milipore, UK). Data were acquired using guavaSoft 2.7 software for Windows and analysed using its 3.1.1. version. Data were presented as median fluorescence intensity indicating the level of marker’s expression (MFI of CD14^+^ and MFI of TLR-2^+^ cells). Doublet and debris exclusion were performed and gating was set based on Fluorescence Minus One (FMO) controls.

### Measurement of TNF-α levels by ELISA

Stored supernatants were used for the analysis of TNF-α concentration by ELISA (TNF alpha (Total) Human ELISA Kit, Invitrogen, UK) following the manufacturer’s instructions. Briefly, the plate was coated with TNF-α specific capture antigen and left overnight. Following subsequent washes, non-specific binding was blocked using blocking buffer after which samples were added and incubated for 2 h at room temperature. Plates were then washed and biotinylated detection antibody was added, after which avidin horseradish peroxidase (avidin-HRP) was used to bind to the biotin on the detection antibody. Tetra-methyl benzidine was then added as the substrate for the HRP enzyme, and the reaction was stopped with the addition of 1M sulfuric acid. Absorbance was read at 495nm using FLUOstar Optima plate reader (BMG Labtech). The data were analysed using MARS Data Analysis Software, and a 4 parameter logistic regression curve was used as the best fit for the data.

### Statistical analyses

Due to the variability between the donors, each was treated as a separate group and Two-Way analysis of variance (ANOVA) followed by Tukey’s posthoc test for multiple comparison were used to examine the differences between the conditions. For each donor, data are presented as mean ± SEM (standard error of the mean) of the replicates of the conditions set up in experiment.

## Supplementary Data

**Figure S1:**
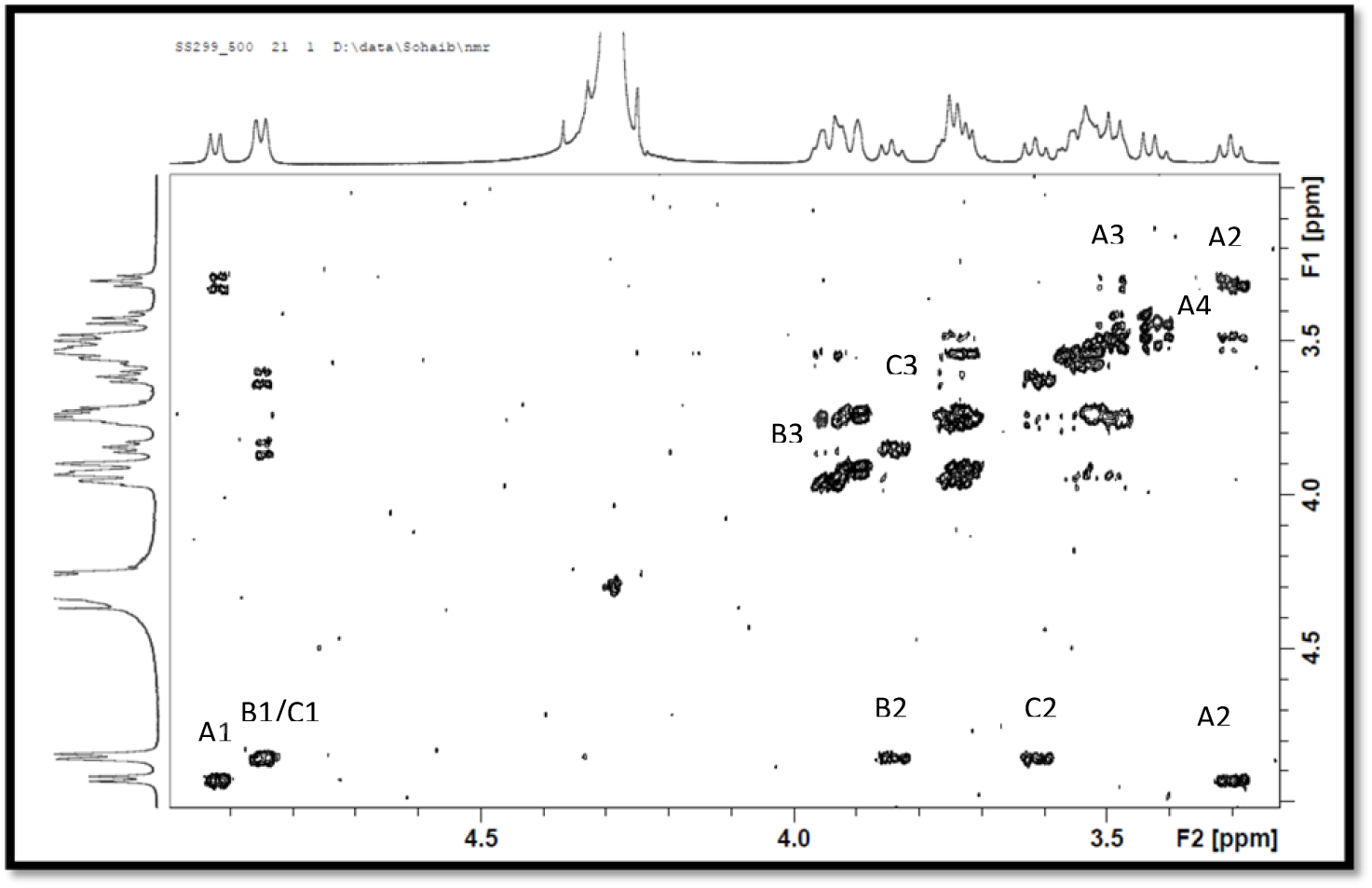
^1^H-^1^H COSY spectrum for the HMw β-glucan recorded in solution in D_2_O (5-10 mg in 0.65 mL) at 70 °C on a Bruker 500MHz spectrometer, Labels (A-C) identify the different monosaccharides and the numbers(1-3) identify the respective proton/carbon.

**Figure S2:**
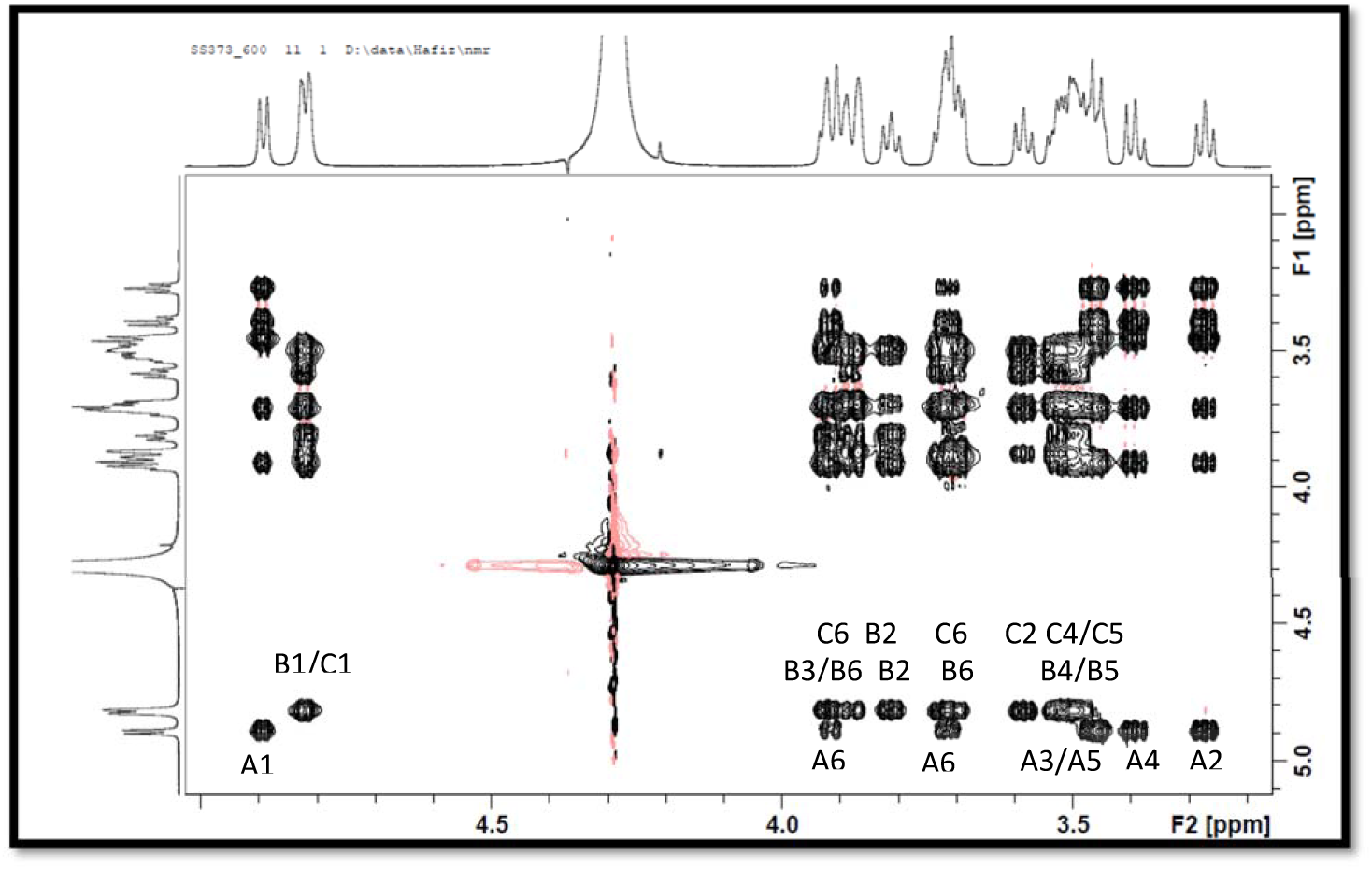
^1^H-^1^H TOCSY spectrum for the HMw β-glucan recorded in solution in D_2_O (5-10 mg in 0.65 mL) at 70 °C on a Bruker 600MHz spectrometer, Labels (A-C) identify the different monosaccharides and the numbers (1-6) identify the respective proton/carbon.

**Figure S3:**
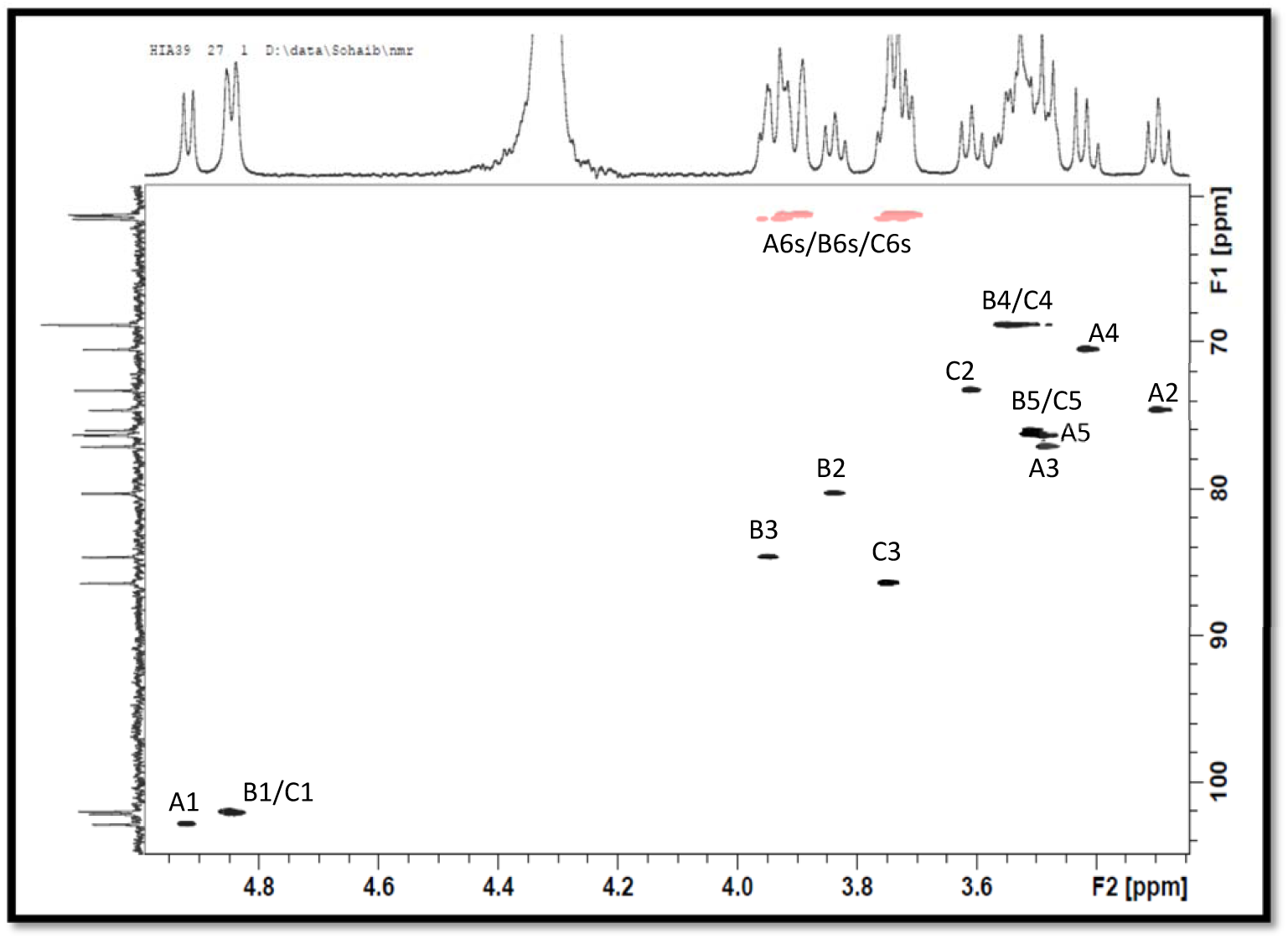
^1^H-^1^H HSQC spectrum for the HMw β-glucan recorded in solution in D_2_O (5-10 mg in 0.65 mL) at 70 °C on a Bruker 500MHz spectrometer, Labels (A-C) identify the different monosaccharides and the numbers (1-6) identify the respective proton/carbon.

**Figure S4:**
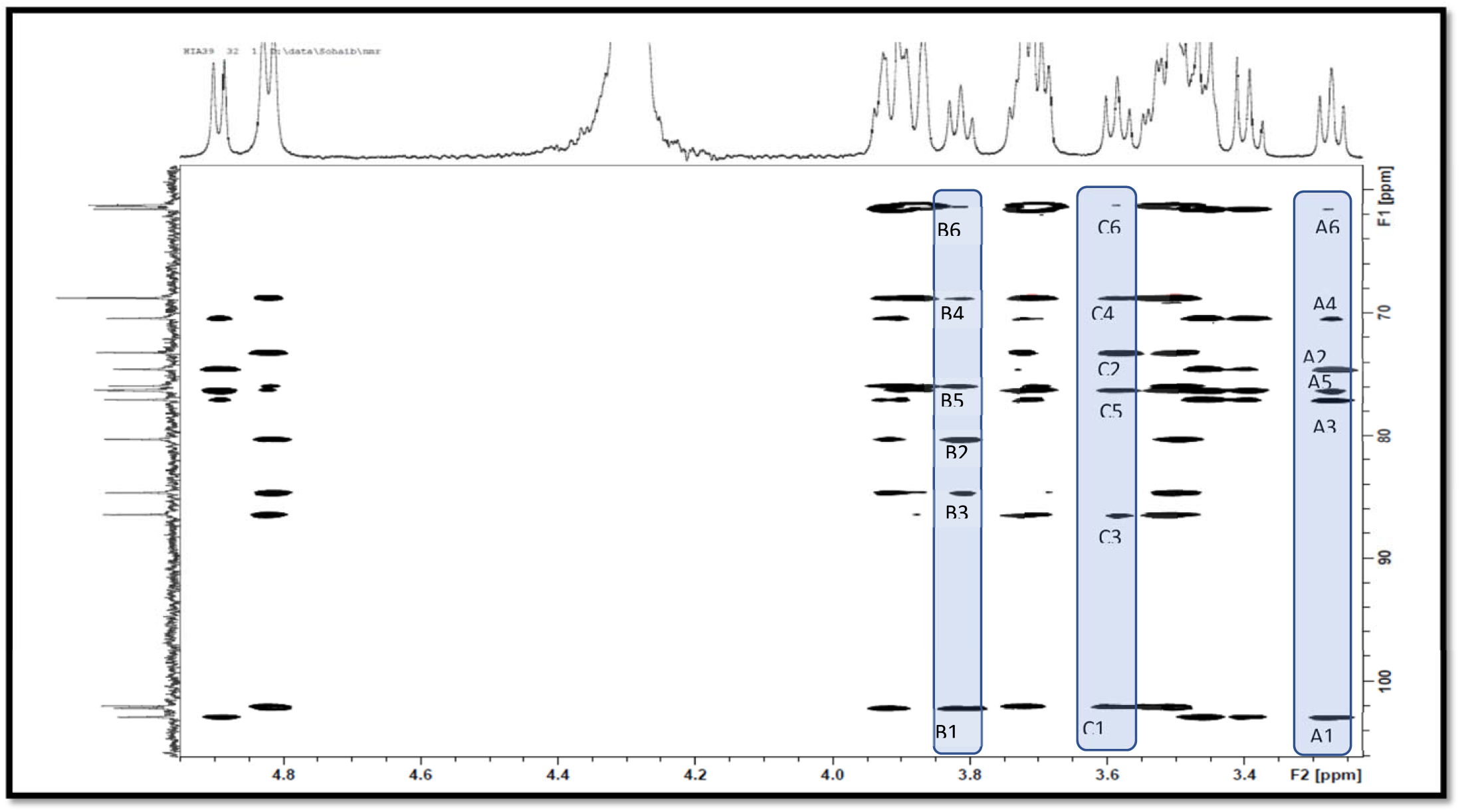
^13^C-^1^H HSQC-TOCSY spectrum for the HMw β-glucan recorded in solution in D_2_O (5-10 mg in 0.65 mL) at 70 °C on a Bruker 400MHz spectrometer, Labels (A-C) identify the different monosaccharides and the numbers (1-6) identify the respective proton/carbon.

**Figure S5:**
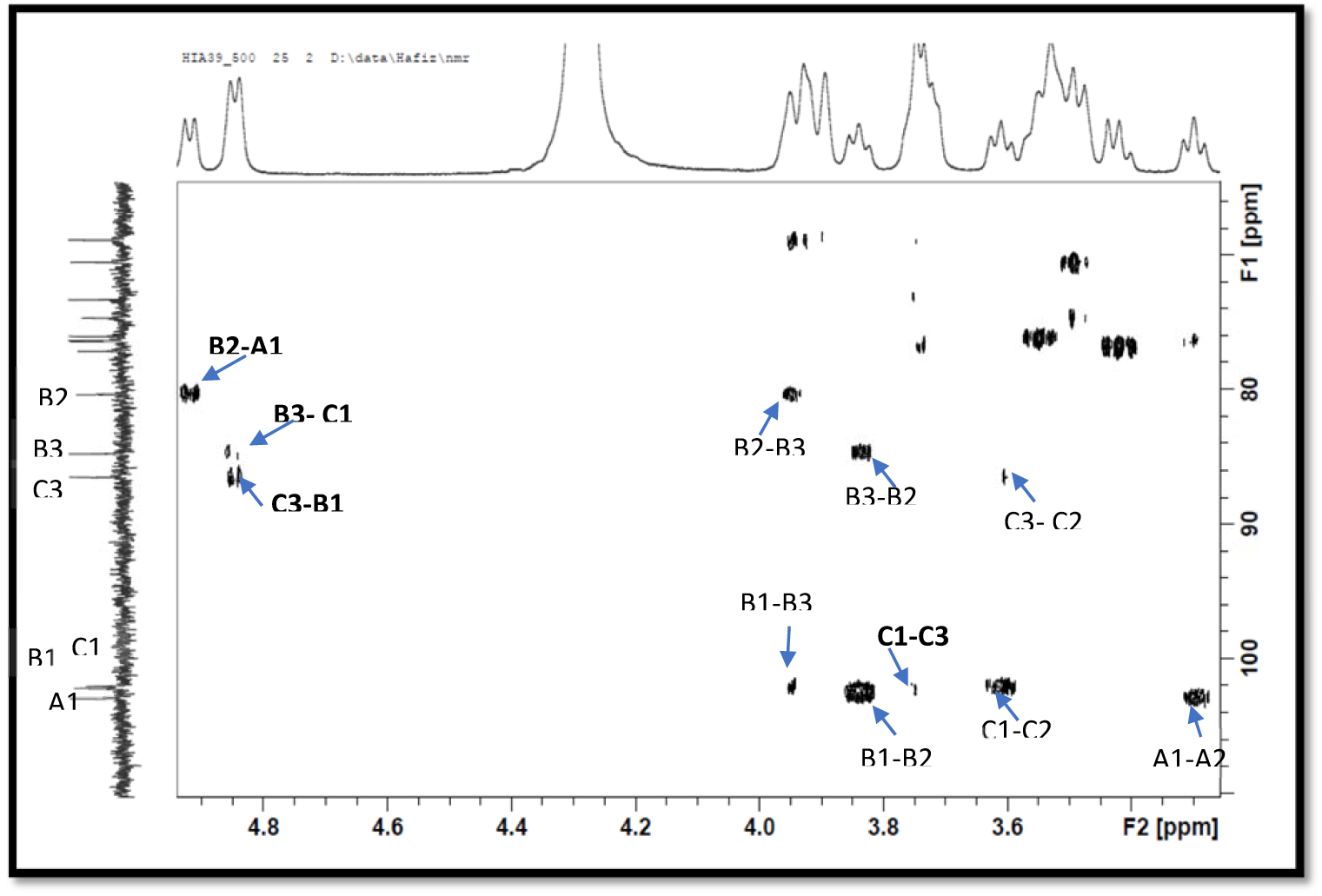
^13^C-^1^H HSQC-HMBC spectrum for the HMw β-glucan recorded in solution in D_2_O (5-10 mg in 0.65 mL) at 70 °C on a Bruker 500MHz spectrometer, Labels identify the significant inter and intra residue scalar coupling, those involved in linkages are highlighted in bold; (A-C) identify the different monosaccharides and the numbers (1-6) identify the respective proton/carbon.

**Figure S6:**
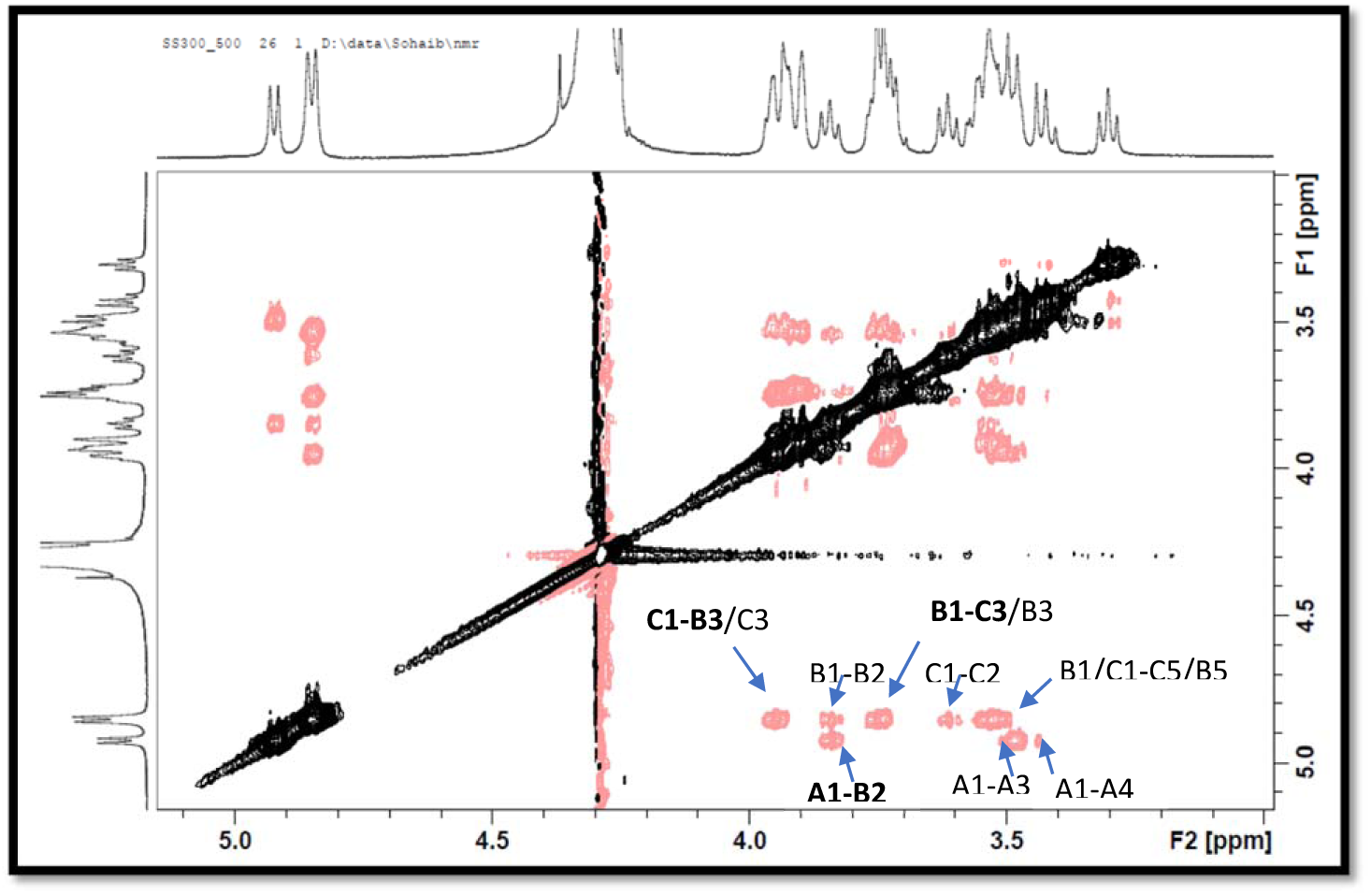
^1^H-^1^H ROESY spectrum for the HMw β-glucan recorded in solution in D_2_O (5-10 mg in 0.65 mL) at 70 °C on a Bruker 500MHz spectrometer, Labels the significant inter and intra residue NOEs-those locating linkages are highlighted in bold involved (A-C) identify the different monosaccharides and the numbers (1-6) identify the respective proton/carbon.

## References

1. Grover S, Sharma VK, Mallapa RH, Batish VK. 2013. Draft genome sequence of *Lactobacillus fermentum* Lf1, an Indian isolate of human gut origin. Genome Announcements 1(6) PMC3828308

2. Jiménez E, Langa S, Martín V, Arroyo R, Martín R, Fernández L, Rodríguez JM. 2010. Complete genome sequence of *Lactobacillus fermentum* CECT 5716, a probiotic strain isolated from human milk. Journal of Bacteriology 192:4800

3. Jayashree S, Pooja S, Pushpanathan M, Vishnu U, Sankarasubramanian J, Rajendhran J, Gunasekaran, P. 2013. Genome sequence of *Lactobacillus fermentum* strain MTCC 8711, a probiotic bacterium isolated from yogurt. Genome Announcements 1(5) PMC3784788

4. Pereira DIA, McCartney AL, Gibson GR. 2003. An in vitro study of the probiotic potential of a bile-salt-hydrolyzing *Lactobacillus fermentum* strain, and determination of its cholesterol-lowering properties. Applied and Environmental Microbiology 69:4743–52

5. Ramos CL, Thorsen L, Schwan RF, Jespersen L. 2013. Strain-specific probiotics properties of *Lactobacillus fermentum, Lactobacillus plantarum* and *Lactobacillus brevis* isolates from Brazilian food products. Food Microbiology 36:22–9

6. Sanni AI, Morlon-Guyot J, Guyot JP. 2002. New efficient amylase-producing strains of *Lactobacillus plantarum* and *L. fermentum* isolated from different Nigerian traditional fermented foods. International Journal of Food Microbiology 72:53–62

7. Verce M, De Vuyst L, Weckx S. 2018. Complete and annotated genome sequence of the sourdough lactic acid bacterium *Lactobacillus fermentum* IMDO 130101. Genome Announcements 6(19) PMC5946041

8. Maldonado J, Cañabate F, Sempere L, Vela F, Sánchez AR, Narbona, E. López-Huertas E, Geerlings A, Valero A. D, Olivares M, Lara-Villoslada F, 2012. Human milk probiotic *lactobacillus fermentum* CECT5716 reduces the incidence of gastrointestinal and upper respiratory tract infections in infants. Journal of Pediatric Gastroenterology and Nutrition 54:55–61

9. Mikelsaar M, Zilmer M. 2009. Lactobacillus fermentum ME-3 - An antimicrobial and antioxidative probiotic. Microbial Ecology in Health and Disease 21:1–27

10. Shi T, Aryantini NP, Uchida K, Urashima T, Fukuda K. 2014. Enhancement of exopolysaccharide production of *Lactobacillus fermentum* TDS030603 by modifying culture conditions. Biosci Microbiota Food Health 33:85–90

11. Ale EC, Perezlindo MJ, Burns P, Tabacman E, Reinheimer JA, Binetti AG. 2016. Exopolysaccharide from *Lactobacillus fermentum Lf2* and its functional characterization as a yogurt additive. Journal of Dairy Research 83:487–92

12. Balzaretti S, Taverniti V, Guglielmetti S, Fiore W, Minuzzo M, Ngo, H. N. Ngere, JB, Sadiq S, Humphreys PN, Laws AP. 2017. A novel rhamnose-rich hetero-exopolysaccharide isolated from *Lactobacillus paracasei DG* activates THP-1 human monocytic cells. Applied and Environmental Microbiology 83

13. Patten DA, Leivers S, Chadha MJ, Maqsood M, Humphreys PN, Laws, AP, Collett A. 2014. The structure and immunomodulatory activity on intestinal epithelial cells of the EPSs isolated from *Lactobacillus helveticus* sp. Rosyjski and Lactobacillus acidophilus sp. 5e2. Carbohydrate Research 384:119–27

14. West MA, Heagy W. 2002. Endotoxin tolerance: A review. Critical Care Medicine 30:S64–S73

15. Biswas SK, Lopez-Collazo E. 2009. Endotoxin tolerance: new mechanisms, molecules and clinical significance. Trends Immunology 30:475–87

16. Pugin J, Heumann ID, Tomasz A, Kravchenko VV, Akamatsu Y, et al. 1994. CD14 is a pattern recognition receptor. Immunity 1:509–16

17. Kawai T, Akira S. 2010. The role of pattern-recognition receptors in innate immunity: update on Toll-like receptors. Nature Immunology 11:373–84

18. Uematsu S, Akira S. 2008. Toll-Like receptors (TLRs) and their ligands. Handbook of Experimental Pharmacology:1–20

19. Watters C, Fleming D, Bishop D, Rumbaugh KP. 2016. Host Responses to Biofilm. Progress in Molecular Biology and Translational Science 142:193–239

20. Wang JH, Doyle M, Manning BJ, Di Wu Q, Blankson S, Redmond HP. 2002. Induction of bacterial lipoprotein tolerance is associated with suppression of toll-like receptor 2 expression. Journal of Biological Chemistry 277:36068–75

21. Labeta MO, Durieux JJ, Spagnoli G, Fernandez N, Wijdenes J, Herrmann R. 1993. CD14 and tolerance to lipopolysaccharide: biochemical and functional analysis. Immunology 80:415–23

22. Rajaiah R, Perkins DJ, Ireland DD, Vogel SN. 2015. CD14 dependence of TLR4 endocytosis and TRIF signaling displays ligand specificity and is dissociable in endotoxin tolerance. Proceedings of the National Academy Science U S A 112:8391–6

23. Lin SM, Frevert CW, Kajikawa O, Wurfel MM, Ballman K, Mongovin, S. Wong VA, Selk A, Martin TR. 2004. Differential regulation of membrane CD14 expression and endotoxin-tolerance in alveolar macrophages. American Journal of Respiratory Cell and Molecular Biology 31:162–70

24. Abbas A, Lichtman AH, Pillai S. 2017. Cellular and Molecular Immunology. Elsevier. 608 pp.

25. Kleiveland CR. 2015. Peripheral Blood Mononuclear Cells. In The Impact of Food Bioactives on Health: in vitro and ex vivo models, ed. Verhoeckx K, Cotter P, Lopez-Exposito I, Kleiveland C, Lea T, Mackie A, Requena T, Swiatecka D, Wichers H. l:161–7. Spinger, Cham (CH). Chapter 15, 161-7.

26. Dueñas-Chasco MT, Rodríguez-Carvajal MA, Mateo PT, Franco-Rodríguez G, Espartero JL, Irastorza-Iribas A, Gil-Serrano AM. 1997. Structural analysis of the exopolysaccharide produced by *Pediococcus damnosus 2.6*. Carbohydrate Research 303:453–8

27. Dueñas-Chasco MT, Rodríguez-Carvajal MA, Tejero-Mateo P, Espartero JL, Irastorza-Iribas A, Gil-Serrano AM. 1998. Structural analysis of the exopolysaccharides produced by *Lactobacillus* spp. G-77. Carbohydrate Research 307:125–33

28. Norian R, Delirezh N, Azadmehr A. 2015. Evaluation of proliferation and cytokines production by mitogen-stimulated bovine peripheral blood mononuclear cells. Veterinary Research Forum 6:265–71

29. Molaee N, Mosayebi G, Pishdadian A, Ejtehadifar M, Ganji A. 2017. Evaluating the Proliferation of human peripheral blood mononuclear cells using MTT Assay. International Journal of Basic Science in Medicine. 2:25–8

30. Laws A, Gu Y, Marshall V. 2001. Biosynthesis, characterisation, and design of bacterial exopolysaccharides from lactic acid bacteria. Biotechnology Advances 19:597–625

31. Ruas-Madiedo P, De Los Reyes-Gavilán CG. 2005. Invited review: Methods for the screening, isolation, and characterization of exopolysaccharides produced by lactic acid bacteria. Journal of Dairy Science 88:843–56

32. Ibarburu I, Soria-Díaz ME, Rodríguez-Carvajal MA, Velasco SE, Tejero-Mateo P, Gil-Serrano A. M, Irastorza A, Dueñas MT. 2007. Growth and exopolysaccharide (EPS) production by *Oenococcus oeni* I4 and structural characterization of their EPSs. Journal of Applied Microbiology 103:477–86

33. Adeyeye A, Jansson PE, Lindberg B, Henrichsen J. 1988. Structural studies of the capsular polysaccharide from *Streptococcus pneumoniae* type 37. Carbohydrate Research 180:295–9

34. Hwang EN, Kang SM, Kim MJ, Lee JW. 2015. Screening of Immune-Active Lactic Acid Bacteria. Korean Journal for Food Science of Animal Resoucesr 35:541–50

35. Ruiz-Bravo A, Jimenez-Valera M, Moreno E, Guerra V, Ramos-Cormenzana A. 2001. Biological response modifier activity of an exopolysaccharide from *Paenibacillus jamilae* CP- Clinical and Diagnostic Laboratory Immunology 8:706–10

36. Bleau C, Monges A, Rashidan K, Laverdure JP, Lacroix M, Van Calsteren MR, Millette M, Savard R, Lamontagne L. 2010. Intermediate chains of exopolysaccharides from *Lactobacillus rhamnosus* RW-9595M increase IL-10 production by macrophages. Journal of Applied Microbiology 108:666–75

37. Hidalgo-Cantabrana C, Nikolic M, Lopez P, Suarez A, Miljkovic M, Kojic M, Margolles A, Golic N, Ruas-Madiedo P. 2014. Exopolysaccharide-producing *Bifidobacterium animalis* subsp. *lactis* strains and their polymers elicit different responses on immune cells from blood and gut associated lymphoid tissue. Anaerobe 26:24–30

38. López P, Monteserína DC, Meimondea M, de los Reyes-Gavilána C, Margollesa A, Sárez A, Ruas-Madiedoa P. 2012. Exopolysaccharide-producing *Bifidobacterium* strains elicit different *in vitro* responses upon interaction with human cells. Food Research International 46

39. Sungur T, Aslim B, Karaaslan C, Aktas B. 2017. Impact of exopolysaccharides (EPSs) of Lactobacillus gasseri strains isolated from human vagina on cervical tumor cells (HeLa). Anaerobe 47:137–44

40. Zhou X, Hong T, Yu Q, Nie S, Gong D, Xiong T, Xie M. 2017. Exopolysaccharides from *Lactobacillus plantarum* NCU116 induce c-Jun dependent Fas/Fasl-mediated apoptosis via TLR2 in mouse intestinal epithelial cancer cells. Scientific Reports 7:14247

41. Patten DA, Leivers S, Chadha MJ, Maqsood M, Humphreys PN, Laws AP, Collett A. 2014. The structure and immunomodulatory activity on intestinal epithelial cells of the EPSs isolated from *Lactobacillus helveticus* sp. *Rosyjski* and *Lactobacillus acidophilus* sp. 5e2. Carbohydrate Research 384:119–27

42. Gao K, Wang C, Liu L, Dou X, Liu J, Yuan, L. Zhang W, Wang Hl. 2017. Immunomodulation and signaling mechanism of *Lactobacillus rhamnosus* GG and its components on porcine intestinal epithelial cells stimulated by lipopolysaccharide. Journal of Microbiology, Immunology and Infection 50:700–13

43. van Bergenhenegouwen J, Plantinga TS, Joosten LA, Netea MG, Folkerts G, Kraneveld AD, Garssen J, Vos AP. 2013. TLR2 & Co: a critical analysis of the complex interactions between TLR2 and coreceptors. Journal of Leukocyte Biology 94:885–902

44. Landmann R, Knopf HP, Link S, Sansano S, Schumann R, Zimmerli W. 1996. Human monocyte CD14 is upregulated by lipopolysaccharide. Infection and Immunity 64:1762–9

45. Dobrovolskaia MA, Vogel SN. 2002. Toll receptors, CD14, and macrophage activation and deactivation by LPS. Microbes and Infection 4:903–14

46. Angus DC, van der Poll T. 2013. Severe sepsis and septic shock. New England Journal of Medicine 369:2063

47. Galdeano CM, Perdigon G. 2006. The probiotic bacterium Lactobacillus casei induces activation of the gut mucosal immune system through innate immunity. Clinical and Vaccine Immunology 13:219–26

48. Zeuthen LH, Fink LN, Frokiaer H. 2008. Toll-like receptor 2 and nucleotide-binding oligomerization domain-2 play divergent roles in the recognition of gut-derived *lactobacilli* and *bifidobacteria* in dendritic cells. Immunology 124:489–502

49. Ale, EC, Perezlindo M J, Pavón Y, Peralta GH, Costa S, Sabbag N, Bergamini C, Reinheimer J A, Binetti A G. (2016a). Technological, rheological and sensory characterizations of a yogurt containing an exopolysaccharide extract from Lactobacillus fermentum Lf2, a new food additive. Food Research International, 90:259–267

50. Dubois M, Gilles K, Hamilton JK, Rebers PA, Smith F. 1951. A colorimetric method for the determination of sugars. Nature 168:167

51. Dubois M, Gilles KA, Hamilton JK, Rebers PA, Smith F. 1956. Colorimetric Method for Determination of Sugars and Related Substances. Analytical Chemistry 28:350–6

52. Gerwig GJ, Kamerling JP, Vliegenthart JFG. 1978. Determination of the d and l configuration of neutral monosaccharides by high-resolution capillary g.l.c. Carbohydrate Research 62:349–57

53. Stellner K, Saito H, Hakomori SI. 1973. Determination of aminosugar linkages in glycolipids by methylation. Aminosugar linkages of ceramide pentasaccharides of rabbit erythrocytes and of Forssman antigen. Archives of Biochemistry and Biophysics 155:464–72

